# High Content Image Analysis for Quantitation of β-arrestin Cell Surface and Endosomal Translocation

**DOI:** 10.64898/2026.11.19.767837

**Authors:** Emma R Carruthers, Caitlin R M Oyagawa, Raewyn C Poulsen, Natasha L Grimsey

## Abstract

The various roles of β-arrestins 1 and 2 in mediating and regulating signaling via G protein-coupled receptors (GPCRs) and other receptors/channels, as well as being mediators of non-canonical signaling and agonist “bias”, make them frequent subjects of contemporary signaling research. Their characteristic subcellular translocation upon receptor activation – changing from diffusely distributed in the cytoplasm (and nucleus for β-arrestin 1) to concentrated at the plasma membrane, clustering in pits, then endocytosing into vesicles – can be leveraged to monitor activation. Proximity-based assays with fluorescent or luminescent readouts from protein complementation or resonance energy transfer are popular quantitative options, but require the β-arrestin, and often the receptor, to be modified with relatively large tags and are typically performed with overexpression. Translocation of untagged β-arrestin can be observed qualitatively by immunocytochemistry, but few options for quantitation of the associated images are available. Here, we share a protocol pairing automated high-throughput microscopy with high-content image analysis in MetaXpress^®^/MetaMorph^®^ (Molecular Devices) to quantitate membrane-like versus punctate vesicle-like staining and thereby facilitate objective analysis of untagged β-arrestin translocation. Although the specific implementation steps and analysis routines are particular to MetaXpress/Metamorph, the principles are applicable to other software packages. The method is demonstrated for β-arrestin 2 translocation following Parathyroid Hormone Receptor 1 (PTH1R) and Vasopressin Receptor 2 (V2R) activation, with and without barbadin (an inhibitor of β-arrestin internalisation).

## 1. Introduction

β-arrestins 1 and 2 are intracellular adaptor proteins that regulate the signaling and subcellular localization of G protein-coupled receptors (GPCRs) and other types of receptors and channels, as well as acting as signaling mediators themselves (Kee et al., 2024). When inactive, β-arrestins are distributed diffusely in the cytoplasm (and nucleus for β-arrestin 1). Following GPCR activation and cytoplasmic domain phosphorylation, β-arrestins are typically observed to translocate to cell surface-localized GPCRs, resulting in a clear concentration at the plasma membrane within a few minutes and acting to reinforce receptor desensitisation. Subsequently, “Class B” GPCRs and β-arrestins bind each other tightly and are co-internalized into early endosomes, from which β-arrestins can scaffold signaling events and latterly either remain associated with a receptor or other scaffolded proteins in subsequent vesicular trafficking steps, or dissociate and return to a diffuse cytoplasmic distribution. Conversely, β-arrestins dissociate from weakly-bound “Class A” GPCRs prior to internalisation, though can still cluster into pits at the plasma membrane and may still have effects on signal transduction via ongoing conformational changes or downstream protein-protein interactions (Peterson & Luttrell, 2017; Tóth et al., 2024).

The characteristic translocation of β-arrestin from the cytoplasm to membrane is widely used in studying GPCR-initiated signaling – either as a primary signaling readout (particularly useful as a non-G protein-specific measure for screening and/or orphan receptors/ligands with unknown coupling/properties) or to compare with G protein-mediated activation/signaling (for example when studying biased agonism). A range of approaches to evaluate β-arrestin translocation are available. Assays that monitor proximity between β-arrestins and receptors or organelles of interest via protein complementation/modification (e.g. PathHunter® [Van Der Lee et al., 2008], LinkLight ^TM^ [Eishingdrelo et al., 2011], Tango^TM^ [Dogra et al., 2016]) or resonance energy transfer (RET, e.g. Donthamsetti et al. [2015]) are popular and extremely useful tools, with usually good signal:noise and often allowing for kinetic measurements. However, these all call for modification of β-arrestin (and often also the receptor) with addition of a fluorescent protein or enzyme, and typically require overexpression, which pose risks for modifying normal biology. Most such studies monitor interaction between β-arrestin and the receptor of interest, which does not distinguish interactions at the cell surface versus endosomes. An alternative design uses bystander RET or protein complementation to detect proximity to the plasma membrane rather than a specific receptor, which reports on the initial translocation event but not subsequent trafficking (e.g. Donthamsetti et al., 2015; Hauge Pedersen et al., 2021). It is also possible to monitor translocation to other organelle markers such as Rab5 for endosomes, though this has been less widely applied to date (e.g. Carruthers et al., 2026; Daly et al., 2023).

However, these protein complementation/proximity approaches do not allow studying untagged β-arrestin, nor endogenously-expressed β-arrestin/receptor (without specialized gene editing to introduce tags, e.g. White et al. [2020]). Other complementary methods such as intramolecular sensors for β-arrestin conformation provide unique insights but again require β-arrestin to be tagged (Lee et al., 2016). Immunoprecipitation or cell fractionation and western blotting for GPCR-β-arrestin complexes can be performed with endogenously expressed proteins but is laborious and only semi-quantitative (Dwivedi-Agnihotri et al., 2020; Marion et al., 2002; Tréfier et al., 2018). Proximity ligation assay (PLA) offers a quantitative protein proximity method that can be applied to untagged endogenously-expressed proteins, though requires reliable antibodies to both the β-arrestin and receptor of interest that bind in conformations that facilitate the PLA reaction (e.g. Petrillo et al., 2019).

Arrestin translocation can readily be observed by simple immunocytochemistry, but is frequently reported as a qualitative outcome. Puncta can be quantitated relatively easily and this approach has been applied to measuring vesicular translocation, though this was optimized for β-arrestin tagged with fluorescent protein and does not distinguish membrane-localized β-arrestin (Leonard et al., 2013; Oakley et al., 2002).

Here we share a protocol for fluorescence immunocytochemistry with widefield (or confocal) microscopy, then applying high content image analysis for objective quantitative measurement of β-arrestin subcellular redistribution to structures resembling the plasma membrane and endosomes. The analysis is applied objectively and consistently across all images and does not require regions of interest to be drawn manually, nor subjective scoring. This approach is well suited to applications where studying unmodified β-arrestin is desired, or as a complement to proximity-based approaches which may report additional insights from the unbiased nature of the approach (Carruthers et al., 2026), and is feasible for use with endogenously-expressed β-arrestins.

To demonstrate this approach, we quantitated the effect of β-arrestin internalisation inhibitor barbadin (Beautrait et al., 2017) on β-arrestin 2 membrane and endosomal translocation after Parathyroid Hormone Receptor 1 (PTH1R) and Vasopressin Receptor 2 (V2R) expressed in HEK cells.

## 2. Cell Preparation, Stimulation and Immunocytochemistry

### 2.1. Rationale

Successful application of this overall method relies on appropriate preparation of cells and execution of immunocytochemistry that will facilitate subsequent imaging by microscopy with good signal:noise for β-arrestin staining.

Fidelity of the antibody for detecting the β-arrestin(s) of interest is imperative. It is strongly recommended to verify/validate staining with positive/negative controls, for example comparing untransfected cells with those transfected with β-arrestins (in the same species as intended for the final experimental design) and preferably without and with stimulation (either the receptor of interest or a positive control). In our experience, manufacturer claims of suitability for detecting β-arrestin by immunocytochemistry and/or β-arrestin 1/2 selectivity are not always consistent with empiric findings (Table 1, Figure 1). Interestingly, one antibody we tested (“A-1”) was selective for human β-arrestin 1 (despite being marketed as non-selective), but did not report translocation upon V2R stimulation, raising the possibility that the antibody might detect an inactive β-arrestin 1 conformation or post-translational modification. We also observed changes in performance of one antibody (“H-9”) between lots, even as a mouse monoclonal antibody, reinforcing the importance of vigilance even when using a previously published/validated antibody. We selected antibody “BD” for our experiments, which detected both human β-arrestin 1 and 2 with good signal:noise and clear alteration in distribution upon stimulation with an appropriate GPCR agonist.

**Table 1.**
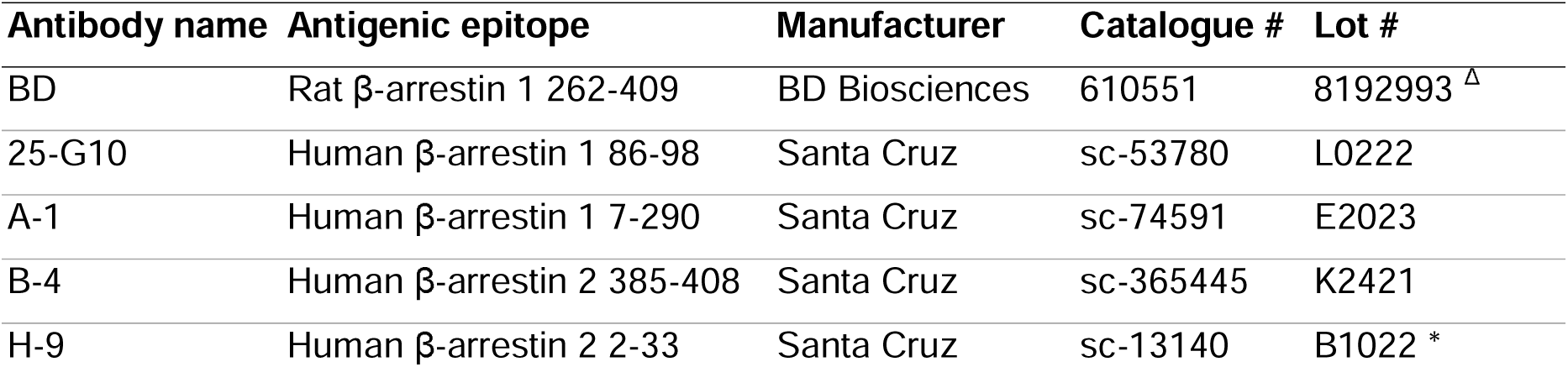
β**-arrestin antibodies tested.** All antibodies raised in mouse. All IgG_1_, except A-1 and B-4 are IgG_2a_. All tested at 1:400 dilution. 25-G10 also tested at 1:50 dilution. See also Figure 1. ^Δ^ An older lot of BD, 41427, performed similarly in prior experiments. * An older lot of H-9, E0906, produced β-arrestin 2 staining concurrently with non-specific punctate staining (data not shown; see also Figure 1).

**Figure 1.**
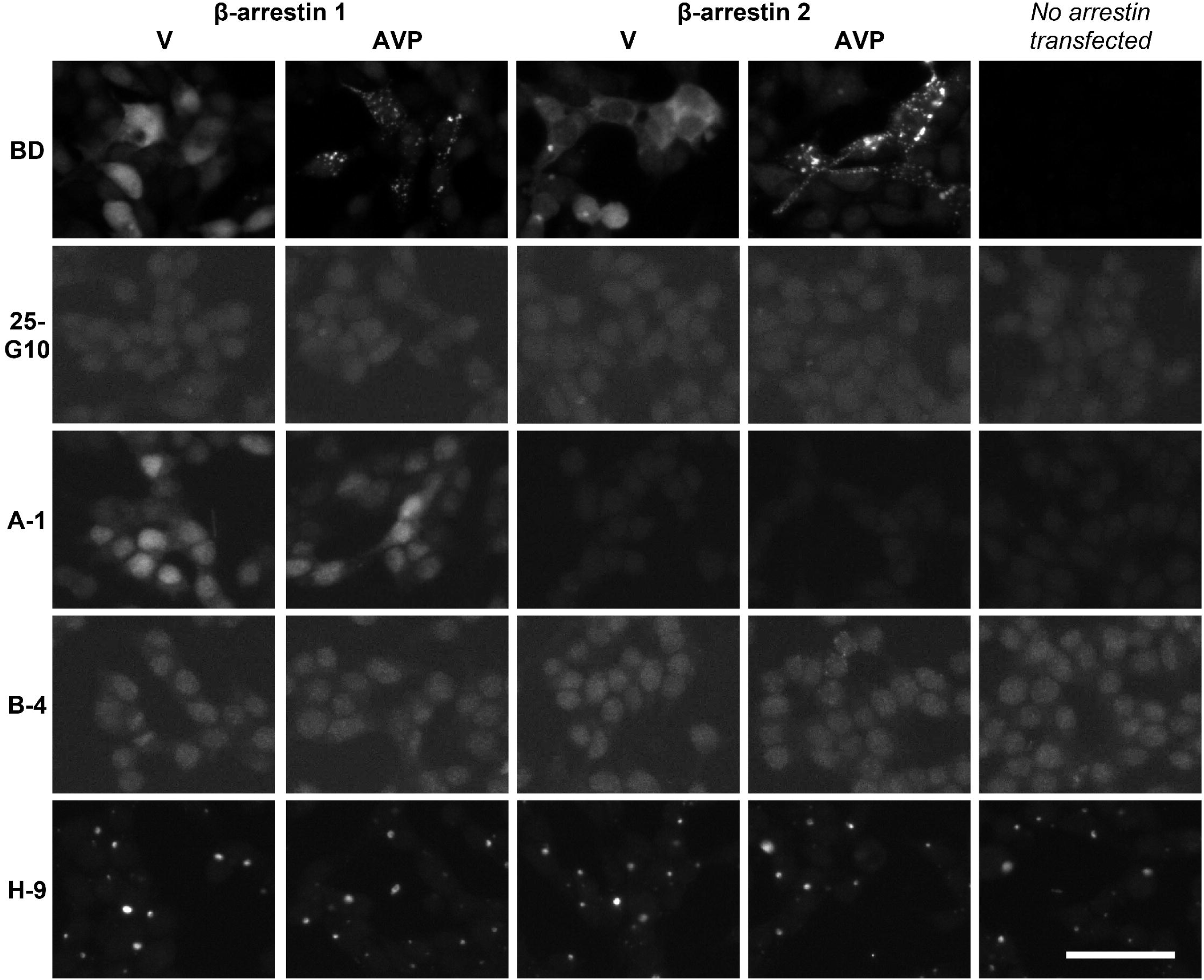
β-arrestin antibody validation. HEK-S cells stably expressing β-arrestin 1 or 2 (or without stable transfection) were transiently transfected with V2R. Cells were stimulated with 1 μM AVP or vehicle (V) for 15 minutes. Antibodies were applied at 1:400 dilution. Within each antibody (rows), images were taken with the same focus and exposure, and are displayed at the same intensity. Antibody “BD” detected both β-arrestin 1 and 2, with staining distribution reflecting translocation following AVP stimulation. “25-G10” yielded no detectable labelling on HEK-β-arrestin 1 or 2 cells. “25-G10” was also tested at a 1:50 dilution, with similar results (data not shown). “A-1” produced positive staining on β-arrestin 1-transfected cells but there was no change in the distribution or intensity of this staining upon AVP stimulation. “B-4” produced low-intensity nuclear staining similarly on all cell lines (including HEK-S without arrestin transfection), but this did not change with stimulation. “H-9” showed bright punctate staining similarly on all cell lines and with no change on stimulation. An older lot of H-9 produced β-arrestin 2 staining concurrently with non-specific punctate staining (data not shown; see also Table 1). Scale bar 50 μm.

The detailed protocols provided here reflect the example data provided in this chapter (Section 4), but many reagent and protocol options are possible.

### 2.2. Materials, Equipment and Reagents

#### 2.2.1 Materials

Cell line or primary cells of interest

- This protocol was developed with adherent cells. The general approach will likely also work with suspension cells with some modifications to preparation for and/or modality of microscopy. It is feasible that a similar approach could work on tissue slices, but more considerable modifications to the overall protocol would be required.
- Cell morphology will impact likelihood of detection of membrane staining; more rounded cells are likely to be more successful than very flat cells. Puncta are expected to be detectable regardless of cell morphology.
- If endogenous β-arrestin is of interest, there will likely be a minimum expression level required to produce sufficient signal, which will also be impacted by the affinity and specificity of the anti-arrestin antibody used.
- If β-arrestin and/or receptor are to be transfected, cells will need to be transfectable.
- We used: HEK-293-S cell line (provided by Professor David Poyner, Aston Research Centre; Wootten et al., 2013).

Plasmids or vector for introducing DNA for expression

- DNA of interest in vector with promoter and/or antibiotic selection gene appropriate for cells of interest.
- Only required if exogenous β-arrestin and/or receptor will be introduced into cells.
- Even if endogenous β-arrestin will be studied, consider using transfected β-arrestins to assist with validating anti-arrestin antibody.
- Optional: Co-expression of plasma membrane and/or organelle markers for validation of analysis parameters and/or β-arrestin localization.
- We used:
  - Human β-arrestin 2 untagged in pcDNA3.1+ (cDNA Resource Center, ARRB200001)
  - Human PTH1R untagged in pcDNA3.1+ (cDNA Resource Center, PTHR100000)
  - Human V2R tagged with 3xHA at the N-terminus in pcDNA3.1+ (cDNA Resource Center, AVR020TN00)
  - Negative control plasmid for antibody validation: Chloramphenicol acetyltransferase (CAT) in pcDNA5/FRT (Thermo Fisher Scientific)

Plates or slides suitable for imaging and compatible with microscope

- For widefield imaging, in our experience standard polystyrene 96-well plates (∼0.8 mm bottom thickness) are adequate, as long as these are of reasonable quality with consistent bottom thickness and no bubbles/scratches in the plastic.
- Imaging may be optimized by using thin-bottom and/or black-wall plates.
- If using a confocal microscope, glass-bottom plates/slides will likely be required.
- We used: Nunc™ MicroWell™ 96-Well, Nunclon Delta-Treated, Flat-Bottom Microplate (Thermo Fisher Scientific, 167008)
- Optional: Gas-permeable plate sealer to prevent evaporation. e.g. AeraSeal Breathable Sealing Film (Excel).

Standard cell culture and immunocytochemistry plasticware and consumables.

#### 2.2.2 Equipment

Standard cell culture facilities – e.g. light microscope, biological safety cabinet, incubator, waterbath, pipettes (preferably multichannel stepper), vacuum aspirator (optional but recommended), metal piece (or card wrapped in foil) that fits inside skirt of culture plate (for cooling cells at end of stimulation).

Standard immunocytochemistry facilities – e.g. fume hood (for fixing cells), rocker.

#### 2.2.3 Reagents

Cell culture maintenance – standard reagents for cell type of interest. We used:

- Dulbecco’s Modified Eagle Medium (DMEM) with high glucose, L-glutamine, sodium pyruvate (Cytiva, SH30243)
- Fetal bovine serum (FBS) (Moregate, New Zealand origin), at 8% in DMEM for cell maintenance (“growth media”)
- Trypsin-EDTA, 0.05% (Thermo Fisher Scientific)
- Poly-D-Lysine (PDL; Sigma-Aldrich); stocks prepared at 1 mg/mL in water
- Phosphate Buffered Saline (PBS; Thermo Fisher Scientific)
- Bovine Serum Albumin (BSA), low endotoxin and low fatty acid (AlbumiNZ)

Transfection (if applicable) – many different reagent options are possible, with optimization/adaptation for cell type. We used:

- Lipofectamine™ 2000 (Thermo Fisher Scientific)
- Opti-MEM™ (Thermo Fisher Scientific)
- PEIMax^®^ (Polysciences) – stock prepared at 1 mg/mL in water, adjusted to pH 7. Stored as single-use aliquots at -20°C.
- Selection antibiotic appropriate for expression plasmid. We used: G418/Geneticin (Gibco).
- Optional: Restriction enzyme suitable for linearizing expression plasmid in a non-functional section of the plasmid. We used: ScaI (Roche).

Stimulations – as applicable for the receptor / research question of interest. Be sure to compare stimuli with matched vehicle control as appropriate. We used:

- hPTH(1-34) (teriparatide acetate; Chemenu). Stock prepared at 1 mM in 0.1% BSA/PBS, stored in aliquots at -80°C allowing no more than three freeze-thaws.
- 8-L-arginine-vasopressin (AVP; Cayman). Stock prepared at 1 mM in 0.1% BSA/PBS, stored in aliquots at -80°C allowing no more than three freeze-thaws.
- Barbadin (β-arrestin-AP2 interaction inhibitor; MedChemExpress). Stock prepared at 10 mM in anhydrous DMSO, stored single-use aliquots at -80°C.
- Plus-One Repel-Silane (GE Healthcare) used to coat plasticware for preparing barbadin dilutions.

Immunocytochemistry – many different reagent options are possible. We used:

- Paraformaldehyde (PFA) 4% in phosphate buffer (Electron Microscopy Sciences). Stored in aliquots at -20°C. Working aliquot at 4°C for up to 1 month.
- Methanol 90% in MilliQ water, ice-cold.
- PBS-T: PBS with 0.2% Triton X-100 (Sigma-Aldrich)
- PBS-T-M: PBS-T with 0.4 mg/mL Merthiolate (BDH)
- Immunobuffer: PBS-T-M with 1% normal goat serum (Sigma-Aldrich)
- Primary anti-β-arrestin antibody: BD Biosciences 610551 “clone 10”, mouse monoclonal, raised against rat β-arrestin 1 262-409. Used at 1:400 in immunobuffer. See Table 1 and Figure 1 for other antibodies tested.
- Secondary antibody: goat anti-mouse Alexa-488 highly cross-adsorbed (Thermo Fisher Scientific, A11029), used at 1:400 in immunobuffer.
- Nuclear dye: Hoescht 33258 (Applichem), stock prepared at 4mg/mL in water, used at 1:500 in PBS-T.
- Optional: antibody or stain for plasma membrane and/or organelle markers for validation of analysis parameters and/or β-arrestin localization.

### 2.3. Protocols

#### 2.3.1 Cell transfection (if applicable)

In the example data provided (Section 4), we used HEK-293-S cell lines stably expressing β-arrestin 2, which were transiently transfected with either PTH1R or V2R. We have elsewhere reported use of the analysis method with transient transfection of both β-arrestin and receptor (Carruthers et al., 2026). We anticipate the analysis method will also be suitable for use with untransfected cells endogenously expressing sufficient levels of β-arrestin 1 and/or 2.

The brief protocols provided here are as used for the data shown in Section 4. Many alternative reagents and protocols are possible, with optimization/adaptation for specific cell types and/or plasmids.

##### 2.3.1.1 Stable transfection

1. Plate cells the day before transfection at 220,000 cells per well in growth media in 24 well plates.
2. Dilute Lipofectamine 2000 (after mixing gently) at 2 µL Lipofectamine in 50 µL OptiMEM per well. Flick mix gently but thoroughly.
3. Dilute 0.8 µg plasmid DNA in 50 µL OptiMEM per well. Mix well (flick or vortex).
  a. Also prepare a tube without DNA as a negative control.
  b. Optional: Rate of successful stable integration of the gene of interest may be enhanced by linearizing the plasmid in a non-functional section prior to proceeding to transfection. We have had mixed success with this approach and often prepare stable transfections with and without linearization for comparison. Incubate 0.8 µg plasmid DNA with 1 unit restriction enzyme and appropriate buffer, making up reaction to 10 µL with nuclease-free water. Incubate for 1 hour at 37°C.
4. Combine diluted Lipofectamine and DNA 1:1. Flick mix gently but thoroughly.
5. Incubate for 20 minutes at room temperature.
6. Pipette onto plated cells, dropwise.
7. Approximately 24 hours after transfection, trypsinise cells and transfer to T25 flasks.
8. After overnight incubation, add selection antibiotic. We used: G418 at 550 µg/mL.
9. Monitor cells over subsequent days, with passaging as needed.
10. Once negative controls contain no viable cells, selection antibiotic concentration can be reduced for maintenance. We used G418 at 400 µg/mL.
11. Screen cells for expression by immunocytochemistry and expanded for cryopreservation. If homogenous or a specific expression level is desired, the population can be clonally isolated or enriched for an expression range (e.g. FACS).

##### 2.3.1.2 Transient transfection

1. Plate cells at 40,000 cells in 80 µL per well growth media in a 96-well plate suitable for imaging, treated with PDL (see Section 2.3.2).
  a. This density is suitable for transfection on the same day. If next day transfection is preferred, plating density may need to be reduced.
2. Dilute PEIMax® in serum-free DMEM; 0.36 µL per well (allow for some excess). Vortex.
3. Dilute plasmid DNA in serum-free DMEM; 75 ng DNA per well (allow for some excess). Vortex.
4. Combine diluted PEIMax® and plasmid in equal volumes. Vortex.
5. Incubate mixes for 60 minutes at 19-21°C.
  a. We have found the optimal incubation time to be temperature-sensitive.
6. Pipette 40 µL per well to cells. Assays were conducted following 18-24 hours expression time.

#### 2.3.2 Cell preparation, stimulation and fixation

The brief protocols provided here are as used for the data shown in Section 4. Many alternative reagents and protocols are possible, with optimization/adaptation for specific cell types, receptors, or stimulations.

1. General recommendations for experimental design:
  a. Include at least two, preferably three, replicate wells per treatment condition.
  b. Avoid using edge wells of plates (greater evaporation from edge wells is frequently observed); instead, fill with PBS or media. Alternatively, use a gas-permeable plate sealer to promote consistent evaporation across plate.
  c. To reduce risk of plate position effects impacting results, avoid systematic positioning of conditions within the plate.
    i. Re-arrange (e.g. randomize) the plate map between independent experiments.
    ii. For large experiments, and/or while establishing consistency and reproducibility, consider including more than one positive and negative control treatment condition in different areas of the plate.
  d. If performing a timecourse experiment we recommend staggering addition of stimulations and fixing the entire plate at once, rather than staggering fixation.
2. Treat culture plate if required for cell type of interest.
  a. Incubate PDL (0.05 mg/mL in PBS) for at least 1 hour (up to overnight) at 37degC. Wash with PBS. Remove.
3. Plate cells as appropriate for cell type and experimental design, aiming for cells to be ∼80% confluent by the time of the assay. If transiently transfecting, follow Section 2.3.1.2.
4. Treat/stimulate cells as appropriate for the experimental design. It is often most efficient, and reduces cell disturbance, to make drugs up at higher than the final intended concentration then add to existing media on wells to dilute. Consider how efficiently solutions will mix upon addition and solubility; we prefer to prepare and add drugs at no more than 5x the final intended concentration. Accuracy is important at all steps.
  a. Aspirate growth media from cells. Equilibrate cells in assay media (DMEM with 1 mg/mL BSA) for 30 minutes. This and later incubations at 37°C in 5% CO_2_.
  b. Prepare barbadin dilutions in assay media in silane-coated tubes to prevent barbadin binding to plastic (recommended in personal communication from Professor M Bouvier). Pre-incubate barbadin (or matched vehicle) on cells for 30 minutes. Here, we prepared barbadin at 4x the final intended concentration (indicated on graphs) then added 1:1 with existing media in plates for pre-incubation, i.e. barbadin was pre-incubated at 2x the final intended concentration. Barbadin was diluted to final intended concentration with subsequent addition of receptor ligand.
  c. Prepare ligands in assay media and add to cells for indicated stimulation times. We prepared ligands at 2x the final intended concentration (indicated on graphs) and added 1:1 with existing media in plates.
5. Restrict/slow receptor binding and signaling by placing the plate on an ice-cold metal surface (cools plate more evenly and rapidly than placing directly on ice chips). Proceed to the next step as soon as possible.
6. Fix cells.
  a. Aspirate media. Working in fume hood, incubate with PFA for 10 minutes. Aspirate and wash with PBS two times, then leave the wells in fresh PBS.
  b. Plates can be stored in the fridge at this point.

#### 2.3.3 Immunocytochemistry

The brief protocols provided here are as used for the data shown in Section 4. Many alternative reagents and protocols are possible, with optimization/adaptation for specific cell types, antibodies or dyes.

1. Optional: Methanol “unmask” epitopes.
  a. Importance of this step is antibody-dependent. When testing new antibodies we recommend testing with and without methanol unmasking. We used methanol unmasking for all experiments in this chapter.
  b. Aspirate PBS. With plate on ice, apply 90% ice cold methanol. Incubate at 4°C for 10 minutes. Aspirate and allow to air dry in fume hood for 3 minutes at room temperature. Re-hydrate with PBS-T for 10 minutes at room temperature.
2. Incubate primary, then secondary, antibody.
  a. Start with primary antibody, then repeat steps with secondary antibody.
  b. Centrifuge antibody to pellet any precipitate; 14,000 xg, 10 minutes, 4°C.
  c. Dilute antibody in immunobuffer.
  d. Incubate on cells with rocking either for 3 hours at room temperature, or overnight at 4°C.
  e. Aspirate and wash once with PBS-T at room temperature, with volume at least 3x that used for antibody incubation.
3. Apply nuclear stain.
  a. Dilute Hoechst in PBS-T.
  b. Incubate on cells with rocking for 15-30 minutes.
  c. Aspirate and wash twice with PBS-T at room temperature, with volume at least 3x that used for antibody incubation. Incubate the second wash for at least 10 minutes.
  d. Aspirate and store in PBS-T-M at 4°C.

### 2.4. Safety considerations and standards

Follow standard institutional biological and chemical safety guidelines as appropriate for cell type, transfection/transduction method, drug treatments, chemicals and equipment used.

Permission to create GMOs may be relevant if introducing DNA. Ethical approval may be relevant if using primary cells.

## 3. Microscopy and High Content Image Analysis

### 3.1. Rationale

As noted in the introduction, the characteristic redistribution of β-arrestin after GPCR activation from diffuse in the cytoplasm to concentrated at the plasma membrane, clustering into clathrin-coated pits, and subsequent internalization to endosomes, is amenable to quantification via fluorescence microscopy paired with appropriate algorithms to identify image features.

It is important that microscopy is carried out in an unbiased manner and excellent consistency between conditions. While it would be feasible to use manual microscopy with strict guidelines in place for avoiding operator bias (e.g. blinding, guides for imaging from the same positions in each well), this would be time-consuming and carry high risks of inconsistent imaging (e.g. focusing, bleaching). Automated microscopy, with consistent focusing, exposure, and well positioning, is ideal for this task. We have developed our method using widefield microscopy, but the general approach and analysis steps would also be suitable for confocal microscopy.

The specific high content analysis routine described here was designed in, and uses tools exclusive to, MetaMorph/MetaXpress, which requires a paid license. However, other software packages with comprehensive analysis tools are likely able to achieve similar outcomes following the same principles (e.g. ImageJ). To segment for membrane-like staining, we utilized the MetaXpress/MetaMorph “Angiogensis Tube Formation” module, designed to detect “tube-like” staining with original applicability for analysis of blood vessels. To the remainder of the image, we applied the “Transfluor” module which detects punctate image features, with the option of distinguishing two puncta sizes, and was originally intended for GPCR internalisation analysis. Prior to applying these modules, filters were applied to assist with enhancing features of interest (note measurements are based on staining area rather than intensity). Analysis was paired with nuclei counting to enable normalization to cell count in each image. As well, an optional threshold-based filter was applied to segment and remove image areas affected by brightly-fluorescent artefact (e.g. secondary antibody precipitation).

### 3.2. Materials, Equipment and Reagents

Fluorescence microscope

- Preferably with mechanical automated stage and focusing for optimal consistency of imaging and to avoid operator bias/artefact.
- Either widefield or confocal imaging should be suitable, though one or other modality might be more successful for different cell types/morphologies.
- Objective lens and camera providing sufficient resolution and contrast for discerning features of interest.
- Excitation lasers and/or filters, dichroic mirrors, and emission filters appropriate for nuclear dye and secondary antibod(ies) used.
- We used: ImageXpress^®^ Micro XLS Widefield High-Content Analysis System (Molecular Devices), with Zyla 4.2 sCMOS 16-bit monochrome camera (Andor), 20x 0.45 NA CFI Super Plan Fluor ELWD ADM objective (Nikon), SPECTRA X solid-state light engine (Lumencor), DAPI/FITC dual emission filter cube (Semrock). Resulting images were 2160×2160 px, 721×721 μm, approximately 1500-2000 HEK cells per image.

Image analysis software

- The specific analysis routine described here is for implementation in MetaXpress^®^ or MetaMorph^®^ (Molecular Devices).
- We used: MetaXpress^®^ v6.5.5.559 with modules “Angiogenesis Tube Formation” 9500-0016 and “Transfluor^®^” 9500-0033.
- Analogous analysis routine/principles could likely be achieved in other software, e.g. ImageJ.

Data processing and presentation

- Standard software for formatting data, performing calculations, and graphing.
- We used: Microsoft Excel (Microsoft) and Graphpad Prism (v10, GraphPad Software).

### 3.3. Protocols

#### 3.3.1 Imaging

1. Allow the plate to equilibrate to room temperature. Check there is a consistent volume of buffer in all wells and no bubbles. Clean the bottom of plate with ethanol and lint-free tissue. Remove the lid prior to imaging.
2. Set up microscope. We used 20x for sufficient resolution to readily segment membrane-like and vesicle-like staining. If applicable, set correction collar on objective lens as appropriate for plate thickness.
3. Decide number of sites/fields to capture in each well. We suggest at least four sites, but recommend 9 sites for more thorough sampling. We used: 9 sites per well, 20 μm spacing between images.
4. Select filters as appropriate for the dye/fluorophore(s) used. Specificity of each filter for the intended fluorophore (i.e. lack of bleed-through) should be verified in pilot experiments. We used: DAPI for Hoechst, FITC for Alexa 488.
5. Establish focus parameters as appropriate for the microscope and substrate. We used: Laser autofocus on the plate and well bottom at each site, with z-offsets for optimized for each wavelength of interest.
6. Set exposure times for each dye/fluorophore. We suggest aiming for the maximum pixel intensity in an image with the brightest expected staining to reach approximately two thirds of the camera pixel depth. This provides good dynamic range while avoiding saturation in case any images have brighter staining. However, if there is particularly bright debris, or a wide range of staining intensity (e.g. transient transfection), it may be necessary to saturate images in some parts to be able to detect low intensity staining. In this case, we recommend taking images at two exposures for greater flexibility in later analysis / processing.

#### 3.3.2 Analysis steps

“Threshold” refers to a pixel intensity/brightness value, or “gray level” in MetaXpress/MetaMorph, that acts as a cutoff for segmenting an image into areas containing pixels with gray levels above versus below the threshold.

1. Optional: If bright artefact such as debris or antibody precipitate is evident in some images, a threshold-based mask can be used to exclude affected image areas from analysis. (Note: considerable or systematic artefact might impact the integrity of the data/analysis and further optimization of immunocytochemistry and/or imaging may be required; see also Section 3.3.3 Step 1).
  a. Select a threshold that is above the brightest staining of interest and that encapsulates all (or at least the center) of the bright artefact. Create a binary “debris” mask based on this threshold (Binary Operations: “Binarize”).
  b. Apply the “Dilate” Morphology Filter to expand the debris mask to encapsulate the full area of debris and out-of-focus blur from the debris (the latter is particularly relevant for widefield imaging; we used 60 px). If the entirety of the bright artefact (with associated out-of-focus blur) is not encapsulated in the dilated debris mask, the image should be excluded from later processing/graphing.
  c. For all images of interest (including both staining of interest and nuclei), replace all pixels in the debris mask area with a gray level approximating the background image intensity (Arithmetic: “Maximum” between the original image and the background gray level). Using a gray level at the level of background in the rest of the image, rather than zero, is required for compatibility with later analysis steps which rely on comparison of intensity levels across the image (“local background”) to identify features.
2. Count cells by applying either “Find Spots” or “Count Nuclei” (standalone or from within “Transfluor”) modules to the nuclear staining image.
  a. “Find Spots” is the faster algorithm and works well for cell types with roughly spherical nuclei with relatively consistent size and intensity. User selects spot size (diameter; we used 30 px) and intensity cutoff (arbitrary units, established/optimized by testing).
  b. “Count Nuclei” is advantageous for cell types with heterogeneous nuclei shapes/sizes. User selects nuclei size minimum and maximum (measured from the image using region tools) and intensity cutoff (in gray values; can be estimated from image then optimized by testing). We typically find the “Standard” algorithm produces more appropriate segmentation than the “Fast” algorithm.
3. Measure overall image area with β-arrestin staining (including diffuse, membrane-like, and punctate) via intensity threshold.
  a. Select a threshold level that segments staining from background. One threshold is selected for the entire experiment and is best judged by comparison to negative controls (e.g. non-transfected cells, no-primary-antibody control) and/or boundary between distinct patterns correlating to cell bodies versus diffuse signals between cells.
  b. The pixel area within the threshold for each image is recorded and can be converted to μm^2^ via the image calibration (if known).
  c. Optional: The average and/or integrated (summed) pixel gray values within the threshold area can also be recorded and may be relevant for checking for consistency of overall β-arrestin expression/staining. In processing/graphing, output values should be background-corrected by subtracting the selected threshold from the average gray value, or subtracting the threshold multiplied by pixel area from integrated gray value. Background-corrected integrated gray value should usually be divided by cell count to account for any fluctuation in cell density between images.
4. Optional: Process β-arrestin image to bring staining intensities between cells into closer alignment. This step is particularly relevant in experiments with markedly different staining intensities between cells (e.g. transiently-expressed or mixed population of stably-expressed β-arrestin) to support subsequent membrane and puncta segmentation that is facilitated by features within an image having relatively consistent intensities. This step is unlikely to be required for clonal stably-expressed or endogenously-expressed β-arrestin.
  a. Create a fast Fourier transform (“FFT”) with “Blur” filter at 90% sensitivity of the β-arrestin image to create a blurred version.
  b. Divide the FFT image by two (Arithmetic: “Divide”), then subtract this from the original image (Arithmetic: “Subtract”). Use the blur-subtracted image for subsequent membrane-like and puncta detection.
5. Detect membrane-like β-arrestin staining by applying filters to enhance feature detection then measuring with the “Angiogenesis Tube Formation” module.
  a. Apply “Unsharp Mask” then “Median” filters from the “Basic Filters” options. These assist in enhancing contrast for membrane-like staining. Settings we used: Unsharp Mask – Filter width 5, Scaling factor 0.9, Result scale 3; Median – Filter width 5, Filter height 5, Sub-sample ratio 1.
  b. Use the “Angiogenesis Tube Formation” module (designed to detect areas of tube-like continuous staining) with “Skeletonization” algorithm. Set the approximate minimum and maximum tubule width; we used 0.3-0.6 μm. An intensity cutoff is also required. In our experiments the module detected membrane-like staining as “nodes”, but with different settings may be reported as “tubules” (if both are detected the areas can be summed).
  c. The membrane-like staining area is recorded in a new image as a binary mask (used in Step 6).
  d. The pixel area within the membrane-like staining mask for each image is recorded and can be converted to μm^2^ via the image calibration (if known).
6. Detect punctate β-arrestin staining by removing the membrane-like staining area from the image then measuring with the “Transfluor” module.
  a. Puncta should be measured on the β-arrestin image from prior to Step 5 (i.e. The filters used during Step 5 to enhance membrane-like staining detection should not be applied).
  b. Apply a small dilation to the membrane-like staining mask from Step 5c (Morphology Filters: “Dilate”; we used 4 px, Circle filter shape). Replace all pixels in the mask area with a gray level approximating the background image intensity (Arithmetic: “Subtract” the dilated membrane-like staining mask from the original image, then “Maximum” between the new image and the background gray level). This avoids the potential for bright membrane-like staining to be re-detected as puncta.
    i. If optional Step 4 (subtract blurred image) was NOT applied, use the background intensity selected in Step 3 (measuring overall β-arrestin staining area) as the “gray level approximating the background image intensity”.
    ii. If optional Step 4 WAS applied, use the background intensity selected in Step 3 multiplied by 0.48 as the “gray level approximating the background image intensity”.
  c. Use the “Transfluor” module to detect spot-like image features. This module allows distinction of smaller puncta (“Pits”, we used 1-2 μm approximate width) versus larger puncta (“Vesicles”, we used 2-3 μm approximate width). Note the distinction between smaller and larger puncta is essentially arbitrary unless further validation is undertaken (see also Section 5). An intensity cutoff is also required. We typically find the “Standard” algorithm produces more appropriate segmentation than the “Fast” algorithm.
  d. The pixel area detected as smaller and larger puncta in each image is recorded and can be converted to μm^2^ via the image calibration (if known).

#### 3.3.3 Analysis implementation and procedure

1. Inspect images for consistency (e.g. focus, background uniformity) and debris/artefact.
  a. Any such issues should be rare and affect random sites/wells. If frequent, or if any systematic artefact is present (e.g. affecting a particular area of the plate, gradient effect), further optimization of imaging and/or immunocytochemistry are likely required.
  b. Images that are out-of-focus or have debris or imaging artefact at a similar intensity level as the staining of interest should excluded from final analysis. Usually analysis will still be performed but the data excluded from later processing/graphing. Selection of images for exclusion should follow objective criteria and be undertaken without reference to the plate map / treatments to reduce risk of investigator bias.
  c. A “montage” overview of all images from the plate is useful and usually sufficient for this assessment. Note the MetaXpress “Review Plate Data” in-built montage overview tool is based on low-resolution image thumbnails which we find is not sufficient; in our implementation we use a custom journal to generate a montage at 20% of the full image size for this purpose.
2. Inspect images for phenotypic features of interest and identify exemplar representative images.
  a. β-arrestins are generally expected to have a diffuse cytoplasmic (and nuclear for β-arrestin 1) distribution in unstimulated cells, translocate to the plasma membrane with short GPCR stimulation, and increasingly punctate staining with prolonged GPCR stimulation.
  b. If no membrane-like staining is qualitatively evident in the overall experiment, it is unlikely valid nor relevant to attempt membrane analysis. To assist with identifying membrane-like staining and optimizing analysis, barbadin may be a useful control as this prevents β-arrestin internalisation and tends to enhance accumulation of β-arrestin at the plasma membrane (see also Section 4).
  c. Identify at least one exemplar image for each of the three types of staining of interest that represents the features.
3. Test and optimize analysis parameters using the exemplar representative image set.
  a. Test and optimize parameters for each analysis step in turn.
  b. “Optimization” aims to find parameters that will produce full detection of the feature of interest in positive control images, without detection of additional unintended image areas, and minimal detection in negative control images. In practice, it is generally not possible to find parameters that will produce perfect segmentation. Figure 2 provides an example of typical segmentation.
  c. We find it helpful to find approximate parameters qualitatively, then run analysis with a few parameter settings on the representative image set to gain a quantitative impression of the optimal setting (and/or to what degree the quantitative window is altered by the setting adjustment). It is still important qualitatively inspect the analysis outcome for the selected parameters to ensure an apparent improvement in quantitative window is not due to artefactual detection of an unintended image feature.
4. Apply analysis to all images in the experiment.
  a. MetaXpress/MetaMorph uses “journals” to record analysis steps and automate application to sets of images. In our implementation, a setup journal prompts the user to select analysis options and enter relevant settings and parameters, which are then saved in local memory. These settings are then the basis for either running the analysis on the full plate, or as the starting point for the next time the analysis setup is run (e.g. if testing multiple setting options during parameter optimization).
  b. Data are either recorded directly in a spreadsheet or text file, or saved into the MetaXpress database for later export.
  c. We can provide our MetaMorph/MetaXpress “journal” files on request. Editing will likely be required for compatibility with individual installations or MetaMorph/MetaXpress versions. Note that the “Angiogenesis Tube Formation” and “Transfluor” modules require specific licenses additional to the base analysis software.

**Figure 2.**
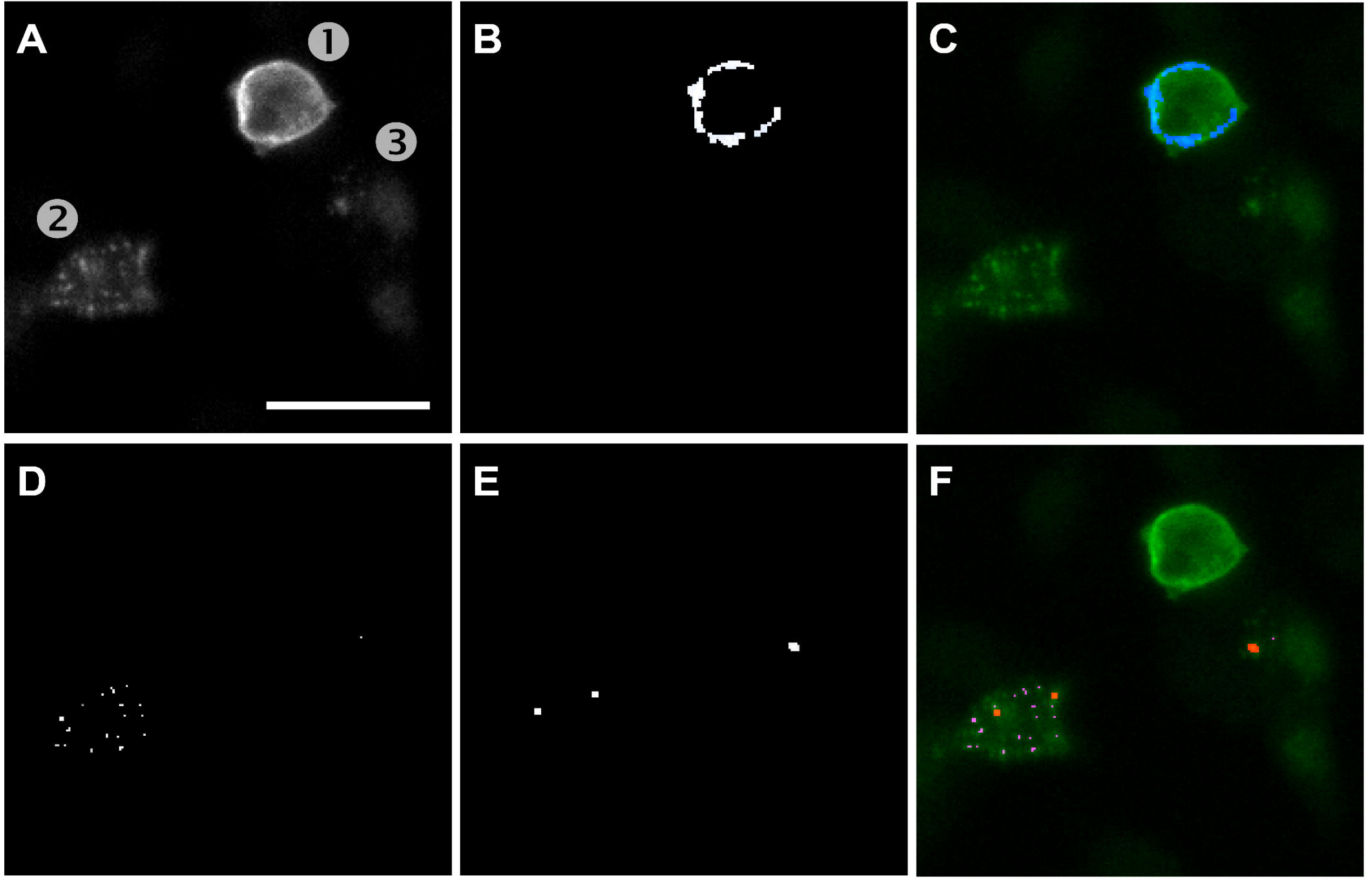
Demonstration of image analysis algorithm segmentation. Representative image of HEK-293 cells stably transfected with β-arrestin 2 and transiently transfected with V2R, stimulated with 1 μM AVP for 15 minutes in the presence of 50 μM barbadin, which produces a mixture of membrane-localized and punctate arrestin staining (quantified results in Figure 4). BD anti-arrestin antibody staining. (A) Cell [1] has predominantly membrane-like staining, [2] has a mixture of small and large puncta, [3] has mainly diffuse arrestin with some low intensity puncta. (B-C) Image in A, indicating area detected by automated analysis as ’membrane-like’ staining (blue). (D-F) Image in A, indicating area detected by automated analysis as ’small’ (E / magenta) versus ’large’ (F / red) puncta. Scale bar 25 μm.

### 3.4. Analysis and statistics

Membrane-like and puncta area measurements can be divided either by the cell count or total β-arrestin area in each image, which adjusts data to account for any variability in cell density. Using the total β-arrestin area may also assist with adjusting for fluctuations in transfection efficiency, if applicable. Cell densities should also be plotted separately to check for any treatment-related alteration in cell density which may impact interpretation. Puncta can be analyzed separately as small versus large puncta areas (and/or the ratio between these), or combined to report total puncta area.

For all measures of interest, calculate the means for all sites in a well (after excluding images with unavoidable artefact, see Section 3.3.2, Step 1), then take the mean from technical replicate wells to produce the means ± SD for a single independent experiment. For statistical analysis, we suggest using these means from three or more independent experiments (without any further normalization); the specific statistical tests and design will depend on the experimental design and research questions.

Due to the imperfect nature of feature segmentation, as well as possible distortion due to the imaging modality (e.g. widefield out-of-focus blur), the absolute areas reported by the analysis should not be considered accurate to the true underlying areas. Furthermore, it is not appropriate to compare membrane versus puncta areas. Therefore, for presentation and commentary, it is likely to be more appropriate to normalize to a control or reference condition and represent results as relative comparisons.

## 4. Results

### 4.1 Barbadin inhibits β-arrestin 2 translocation to endosomes after PTH1R activation

In HEK cells stably expressing β-arrestin 2 and transiently expressing PTH1R, a 15 minute 1 μM hPTH(1-34) stimulation increased β-arrestin 2 membrane-like area, which was not impacted by barbadin at 15 μM or 50 μM (Figure 3A). hPTH(1-34) treatment also induced formation of β-arrestin 2 puncta (Figure 3B). As expected from barbadin’s ability to inhibit β-arrestin internalisation, both concentrations of barbadin inhibited hPTH(1-34)-induced puncta formation.

**Figure 3.**
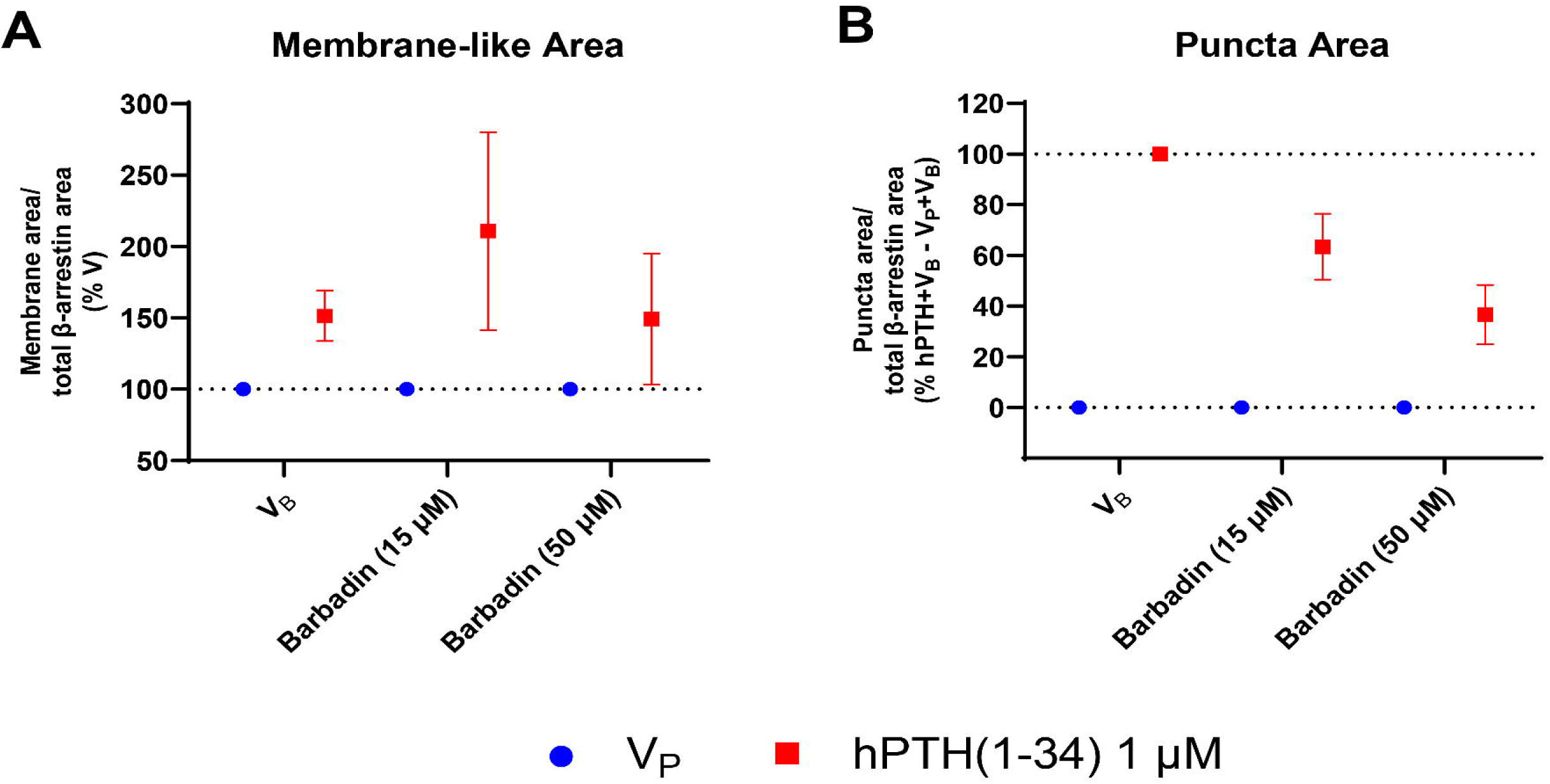
β-arrestin 2 translocation in response to hPTH(1-34) with and without barbadin in HEK cells stably expressing β-arrestin 2 and transiently expressing PTH1R. Cells were pre-incubated with barbadin (15 or 50 µM as indicated) or vehicle (V_B_) then stimulated with hPTH(1-34) (1 µM) or vehicle (V_P_) for 15 minutes. β-arrestin 2 translocation was assessed by ICC with BD anti-β-arrestin and high-content image analysis for membrane-like staining (A) or total puncta (B) area. Data presented as means ± SEM of two independent experiments. Normalized to V_P_+V_B_ (100%) (A), or V_P_+V_B_ (0%) and hPTH(1-34)+V_B_ (100%) (B).

### 4.2 Barbadin enhances β-arrestin 2 translocation to plasma membrane and inhibits translocation to endosomes after V2R activation in a concentration-dependent manner

In HEK cells stably expressing β-arrestin 2 and transiently expressing V2R, AVP 1 μM induced an increase in β-arrestin 2 membrane-like staining, which was enhanced by barbadin at high concentrations (50 μM and 100 μM; Figure 4A). AVP 1 μM also induced an increase in β-arrestin 2 puncta, which was inhibited by barbadin in a concentration-dependent manner (Figure 4B).

**Figure 4.**
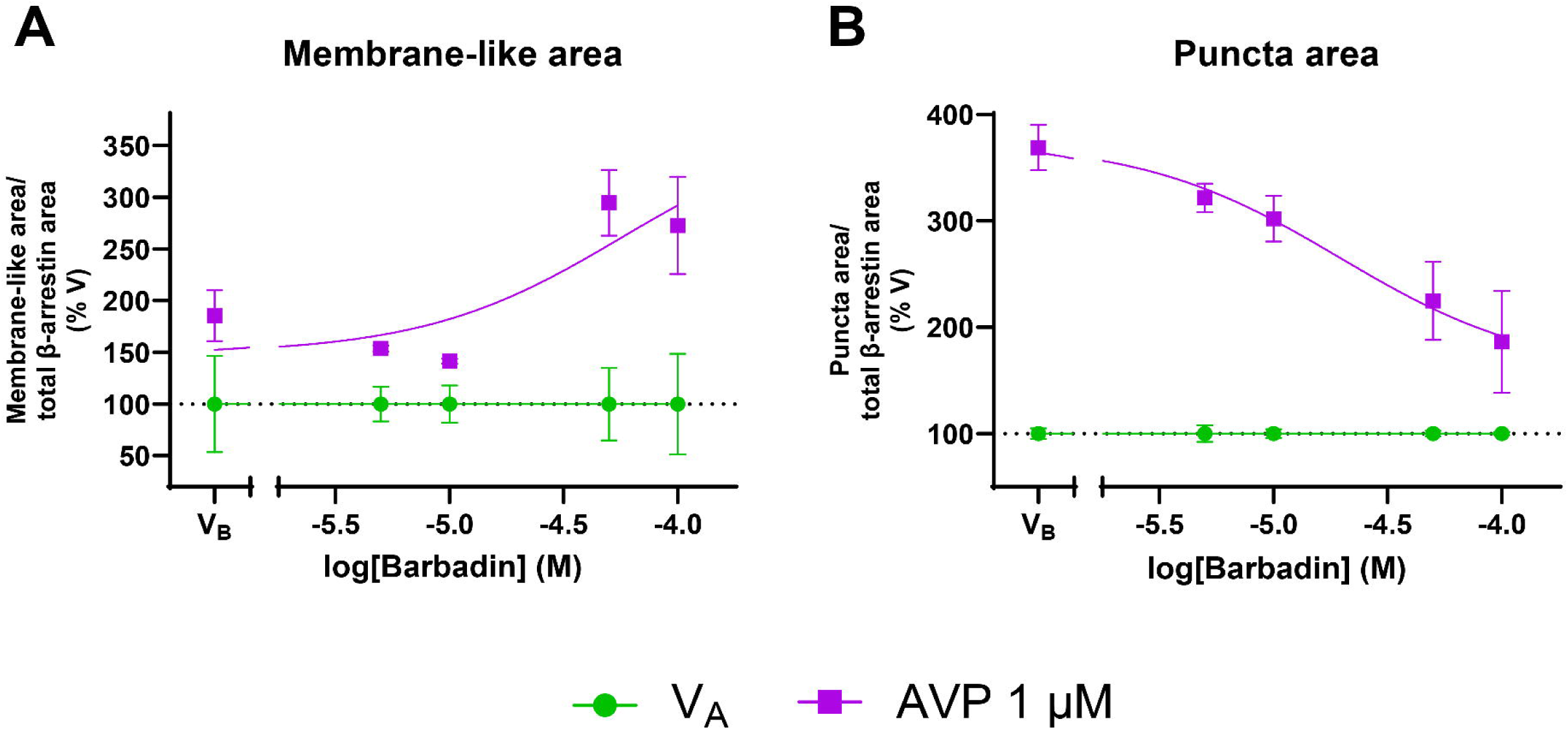
Barbadin concentration response effect on AVP-stimulated β-arrestin 2 translocation in HEK cells stably expressing β-arrestin 2 and transiently expressing V2R. Cells were pre-incubated with a range of concentrations of barbadin (or vehicle, V_B_) for 30 minutes then stimulated with AVP 1 µM or vehicle (V_A_) for 15 minutes. β-arrestin 2 translocation was assessed by ICC with BD anti-β-arrestin and high-content image analysis for membrane-like staining (A) or total puncta (B) area. Data normalized to vehicle for AVP at each barbadin concentration, and presented as means ± SD of technical replicates from one independent experiment.

## 5. Pros and cons

| Pros | Cons |
|---|---|
| Approach can be used with untagged or tagged β-arrestin and receptor. | Assaying untagged β-arrestin requires suitable antibody with good signal:noise. There is potential for results to be impacted by sensitivity of antibody to β-arrestin conformation, interactions, or post-translational modifications (may instead be a ‘Pro’ if such impacts are known / informative). |
| Approach is suitable for exogenously-expressed β-arrestin (either stable or transient expression) and potentially suitable for endogenously-expressed β-arrestin. | Relatively high expression and/or an excellent antibody are likely required for robust analysis.<br>Some receptor-arrestin pairings are likely more feasible for study than others; e.g. “Class A” with short-lived interactions are expected to be more challenging to assay than “Class B”. |
| β-arrestin detection is based on simple immunocytochemistry using one β-arrestin antibody, with simple and rapid sample handling. No co-expression of biosensor components is required (as in protein complementation and RET), and no complex sample handling/preparation (e.g. immunoprecipitation / cell fractionation).<br>Ability to detect β-arrestin translocation is not biased/limited by the complementation partner or RET acceptor. | Membrane-like and punctate staining are not validated as truly representative of underlying membrane or endosomal location or type. Further follow-up required for more definitive classification.<br>In comparison with protein complementation or RET, assay is likely to be less sensitive, lower throughput, and does not produce inherent time-resolved output (need to use staggered stimulation/fixing for time points). |
| Imaging is efficient if using an automated microscope. Manual imaging is also possible. Either widefield or confocal microscopy is suitable. | Preferred requirement for automated microscope. Manual microscopy would be time-consuming and operator/systematic bias would need to be controlled. |
| Analysis is straightforward to undertake in MetaXpress/MetaMorph. It is likely possible to apply the principles to other software. | MetaXpress/MetaMorph is licensed (paid) software and specifically requires the Angiogenesis Tube Formation and Transfluor modules. |
| Analysis is applied objectively and consistently across entire experiment allowing robust relative comparison. | Parameters and resulting segmentation are expected to be imperfect and it is typically necessary to balance “optimal” settings within and between exemplar / control images. Absolute area measurements are unlikely to be accurate (especially with widefield microscopy). |
| Membrane β-arrestin translocation is expected to be relevant for both Class A and B GPCR-arrestin interactions, though detection may be more difficult for Class A due to transient nature of the interaction. | Suitability of membrane measurement may depend on cell type/morphology and imaging modality. General feasibility for visualizing/quantifying membrane could be tested with a membrane antibody / stain / fluorescent protein. |
| Puncta measurement is highly transferrable between cell types and likely to be robust for a range of cell morphologies, as well as applicable to both Class A and B GPCR-arrestin interactions. | Puncta detection might be difficult for Class A GPCRs for which β-arrestin interactions are transient at the plasma membrane and are not sustained during GPCR endocytosis, though β-arrestin has been reported to produce clustered structures even without prolonged GPCR interaction which may well be detectable as small puncta / pits (Eichel et al., 2016). |
| Option to distinguish between small and large puncta, which may represent structures of biological interest – e.g. clathrin-coated pits at the plasma membrane versus endocytosed vesicles. | Nature of small versus large puncta is not inherently validated in the assay, and the structure size parameters set in the analysis do not necessarily enable identification (especially if using widefield microscopy). Further follow-up required for more definitive classification. |

## 9. Troubleshooting & Optimization

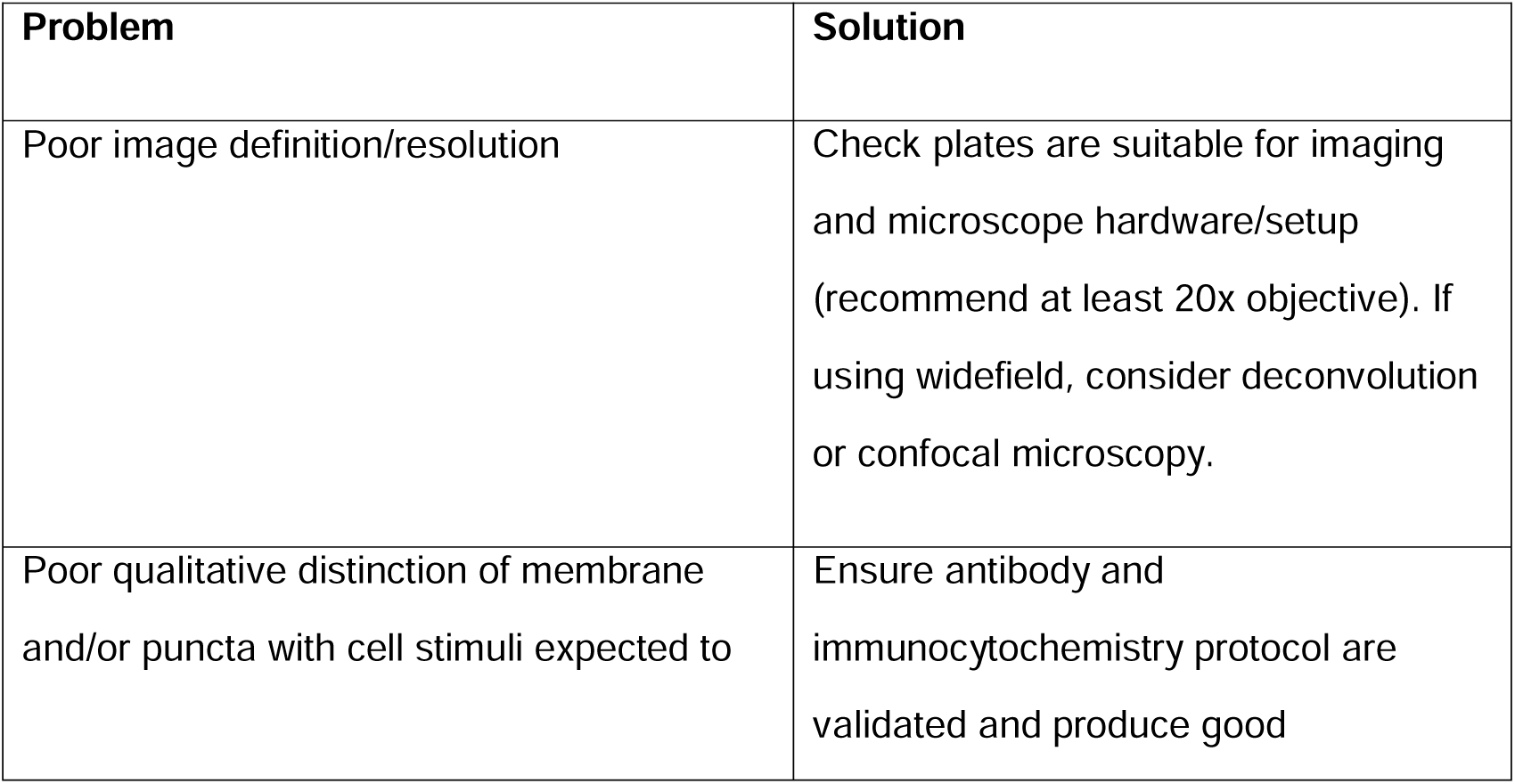
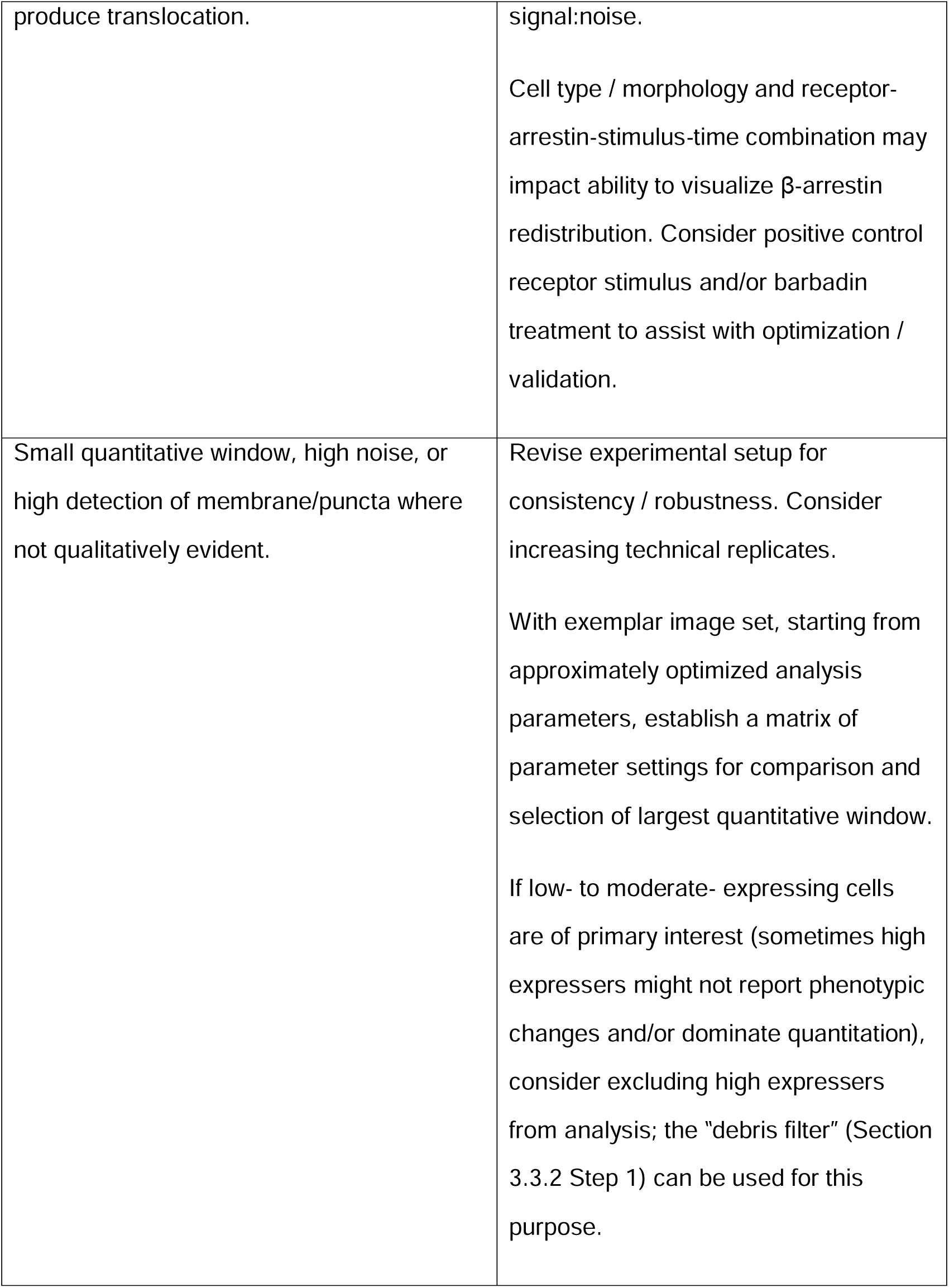

| Problem | Solution |
|---|---|
| Poor image definition/resolution | Check plates are suitable for imaging and microscope hardware/setup (recommend at least 20x objective). If using widefield, consider deconvolution or confocal microscopy. |
| Poor qualitative distinction of membrane and/or puncta with cell stimuli expected to produce translocation. | Ensure antibody and immunocytochemistry protocol are validated and produce good signal:noise.<br>Cell type / morphology and receptor-arrestin-stimulus-time combination may impact ability to visualize β-arrestin redistribution. Consider positive control receptor stimulus and/or barbadin treatment to assist with optimization / validation. |
| Small quantitative window, high noise, or high detection of membrane/puncta where not qualitatively evident. | Revise experimental setup for consistency / robustness. Consider increasing technical replicates.<br>With exemplar image set, starting from approximately optimized analysis parameters, establish a matrix of parameter settings for comparison and selection of largest quantitative window.<br>If low- to moderate- expressing cells are of primary interest (sometimes high expressers might not report phenotypic changes and/or dominate quantitation), consider excluding high expressers from analysis; the “debris filter” (Section 3.3.2 Step 1) can be used for this purpose. |

## Acknowledgements

This research was supported by a Health Research Council of New Zealand Sir Charles Hercus Health Research Fellowship (to NLG; 20-006), a University of Auckland Faculty Research Development Fund Project Grant (to NLG), a University of Auckland Doctoral Scholarship (to ERC), and University of Auckland School of Medical Sciences Graduate Student Research Funding (to ERC and NLG). Microscopy and analysis were performed using instruments in the Biomedical Imaging Research Unit (BIRU) at the University of Auckland.

## Data availability

The data supporting this study’s findings are available from the corresponding author upon reasonable request.

## Author contributions

EC and NLG designed and performed experiments, and analyzed data. CRMO and RCP contributed to study design and interpretation. NLG and EC drafted the manuscript, with contributions and editing from the other authors. All authors approved the submitted version of the manuscript.

## Ethics approval statement

Not applicable.

## Permission to reproduce material from other sources

Not applicable.

